# Evaluation of endothelial nitric oxide synthase gene expression in mice genetically heterozygous in Cx43 (Cx43 + / - mice)

**DOI:** 10.1101/421784

**Authors:** Wilfred Obaalologhi

## Abstract

Gap junctions are proteins made of connexins which are involved in the regulation of vascular function. Deletion of connexins 43 (Cx43) modifies expression of genes known to be involved in the regulation of the vasculature, differentiation and function of vascular cells. Interestingly, mutant mice lacking endothelial nitric oxide synthase (eNOS) gene have been shown to be hypertensive, suggesting that nitric oxide (NO) plays a role in the physiological control of blood pressure. It was therefore hypothesised that the endothelial deletion of Cx43 in the pulmonary vasculature induces endothelial dysfunction and causes eNOS impairment thereby reducing NO biosynthesis, thus leading to vasoconstriction and vascular remodelling which subsequently leads to the development of pulmonary arterial hypertension (PAH). This project was aimed at evaluating eNOS gene expression in mice genetically heterozygous (HET) in Cx43 (Cx43 +/- mice). This was achieved by using lung tissues from four groups of wild type (W/T) and Cx43 +/- (male and female) mice. Ribonucleic acid (RNA) was isolated from the lung tissues using RNA II isolation system and was reverse transcribed to complementary deoxyribonucleic acid (cDNA). End – point polymerase chain reaction (PCR) and real time PCR were used to measure the expression of eNOS gene. eNOS gene expression levels were found to be the same in all four groups of mice tested, with no significant difference. The result therefore suggests that eNOS gene is expressed in mice genetically heterozygous in Cx43 (Cx43 + / -).

## INTRODUCTION

Pulmonary arterial hypertension (PAH) is a fatal progressive disease characterized by persistently elevated pulmonary arterial pressure (PAP), thereby leading to failure of the right ventricle [1]. The disease, in which its aetiology is unknown, is associated with low level of pulmonary NO [2] with clinical manifestations including ventilation–perfusion mismatching, shortness of breath, exercise intolerance, fainting and heart failure in severe cases [3, 4]. From the heart, the pulmonary arteries supply deoxygenated blood to the lungs where gas exchange takes place within the pulmonary capillaries [5]. PAH is a multifactorial disease characterized by constriction and remodelling of the pulmonary vasculature, which is the basis for the persistently high pulmonary pressures [6]. The process of vascular remodelling gives rise to increased and disorganised proliferation of smooth muscle cells (SMCs), pulmonary endothelial cells and fibroblasts, thus resulting in reduction in the number of distal arterial vessels as a consequent of increased muscularization of the arteries, which later progresses and leads to vessel occlusion and then finally, formation of plexiform lesion. Due to increased proliferation, the vessel lumen decreases thereby increasing pulmonary vascular resistance (PVR) and PAP, with a reduction in vascular compliance (pressure = resistance x cardiac output) [7].

Recent data reveal that in PAH patients, the mean resting pulmonary artery pressure rises above 25 mmHg [6] and tends to increase further to 30 mmHg during exercise [8]. Elevated arterial blood pressure is a key risk factor leading to cardiovascular diseases such as right heart failure which is the main cause of mortality In PAH patients [9, 10]. There is poor prognosis and rate of survival from the disease has been reported as only 58% after 3 years on therapy [11]. The disease which occurs more frequently in women than in men presents a remodelled and obliterated pulmonary vasculature, advancing to right ventricular (RV) dysfunction and death [10, 12].

The exact pathogenesis of PAH is not completely understood, however, serotonin, bone morphogenetic protein receptor type 2 and NO pathways have been shown to play major roles in the development of the disease [13, 14]. There is need for novel drugs therapies targeting both vasoconstriction and remodelling, which can only be achieved with known pathogenesis [6]. PAH, which represents group one within the pulmonary hypertension WHO clinical classification system can be further classified, based on aetiology, into idiopathic PAH (IPAH), heritable PAH (HPAH), drug and toxin-induced PAH, associated PAH (APAH) and Persistent pulmonary hypertension of the new born (PPHN) [15].

Regulation of vascular tone and blood flow is mediated by NO which is synthesized by eNOS in the endothelial cells of the pulmonary vasculature. When activated, eNOS catalytically reacts L-arginine and oxygen to produce NO which functions via the NO/cGMP pathway to regulate vascular tone by crossing the endothelium to the vascular smooth muscle and activating soluble guanylyl cyclase which produces cyclic guanosine monophosphate (cGMP) which activates protein kinase G (PKG), thereby mediating smooth muscle relaxation [16]. Expectedly, PDE-5 expression was raised in chronic hypoxic rat, which explains some vascular in the pulmonary vessels as PDE −5 degrades cGMP [17]. Recent studies have shown that NO modulates and maintains arterial pressure [18] and serves as a suppressor of SMC proliferation [19]. The NO pathway is tonically active in resistance vessels, reducing the peripheral vascular resistance and therefore blood pressure [20]. Mutant mice lacking eNOS gene have been shown to be hypertensive, supporting a role for physiological control of blood pressure by NO [20] as genetic mutation of eNOS gene results in failure or insufficient synthesis of NO, giving rise to vasoconstriction and thereby resulting to increase in peripheral blood pressure, which could possibly lead to the formation of atheroma [21]. The enzyme arginase regulates NO biosynthesis through its effect on arginine. Consequently, its activity has been shown to be higher in serum of PAH patients than in controls, suggesting that NO is implicated in the pathogenesis of PAH [2]. Arginase II expression level was increased with a reduction in NO synthesis in pulmonary artery endothelial cells extracted from PAH lungs when compared to control cells in vitro [2].

Cell – cell communication via gap junctions is essential in controlling normal vascular function [22, 23]. These cellular communications involve a family of transmembrane proteins called connexins which oligomerize into a hemichannel and eventually pair with a partner hemichannel in an adjacent cell forming gap junctions that allow cell-to-cell coupling between endothelial and SMC [6, 24]. Key connexin family members involved in the vasculature include connexin 37 (Cx37), connexin 40 (Cx40), connexin 43 (Cx43) and connexin 45 (Cx45), and all play significant role in regulation of vascular function such as mediation of vascular tone, vascular growth, angiogenesis, cell development and differentiation [25] and the passage of cGMP and Ca^2+^, thus regulating vascular tone [26].

Studies have demonstrated that Cx43 expression is restricted to the endothelial cells at branch parts of pulmonary arteries [27]. Genetic deletion of Cx43 in mice corresponds with significantly elevated smooth muscle proliferation [28]. Studies on Cx43 expression in animal models appear to be inconsistent as different effects of Cx43 deletions have been reported [29]. While increased neointimal and adventitia formation were observed in Cx43 knockout mice [28] a recent study however revealed a reduced neointimal formation in Cx43 heterozygous mice [30]. Deletion of Cx43 has been shown to modify expression of genes known to be involved in differentiation and function of vascular cells and cell signalling pathways important for the regulation of vasculogenesis and angiogenesis [31]. Since mutant mice lacking eNOS gene has been shown to be hypertensive, supporting a role for physiological control of blood pressure by NO [20], there seem to be interplay between Cx43 and eNOS expression [32].

The aim of this project was to evaluate the eNOS gene expression in mice genetically heterozygous in Cx43 (Cx43 +/- mice). Lung tissues from four groups of wild type and Cx43 +/- (male and female) mice were used. RNA was isolated from the lung tissues using RNA II isolation system and was reverse transcribed to cDNA. To measure the expression of eNOS gene, end – point PCR and real time PCR were used.

## MATERIALS AND METHODS

### Animal model and Tissue preparation

The investigation conforms to the United Kingdom Animal procedures act 1986, and with the Guide for the Care and Use of Laboratory Animals published by the US National Institutes of Health (NIH publication, 8^th^ Edition, 2011). Ethical approval was granted by the University Ethics Committee. Male and female wild type and connexin 43 heterozygous mice were maintained in group cages, on saw dust bedding, and subjected to a 12 h–12 h light/–dark cycle with food and water provided *ad libitum*. In this study, a total of 12 wild type mice and 12 connexin 43 heterozygous mice aged between 20 and 28 weeks were used. Animals were culled using intraperitoneal administration of pentobarbital sodium (60mg/kg i.p.; JM Loveridge plc, Southampton, UK), lung tissues snap frozen and stored at −80°C until further use. The tissue samples were labelled as shown in table 1.

**Table 1.** Labelled lung tissue samples extracted from the 24 mice

| **HET (Cx 43 +/-)**<br>**male** | **W/T male** | **HET (Cx 43 +/-)**<br>**female** | **W/T female** |
| --- | --- | --- | --- |
| 1 | 1 | 1 | 1 |
| 2 | 2 | 2 | 2 |
| 3 | 3 | 3 | 3 |
| 4 | 4 | 4 | 4 |
| 5 | 5 | 5 | 5 |
| 6 | 6 | 6 | 6 |

### Extraction of RNA using Nucleospin RNA II isolation system

Total RNA was isolated from the mice lung tissues using NucleoSpin^®^ RNA II kit supplied by Macherey-Nagel GmbH & Co. KG, Germany. In proceeding with the RNA extraction from the lung tissues, first, 70% ethanol was prepared from 100% ethanol stock solution. Using the mathematical equation M_1_V_1_ = M_2_V_2_, the volume of water added to ethanol to make it 70% concentrated was obtained by cross multiplying the equation. V1 = volume of stock solution needed = M_2_V_2_ / M_1_ = (70 x 100) / 100 = 70ml. Therefore, 70ml stock ethanol was diluted with 30ml of water, concentrating it to 70%. Furthermore, a DNAse reaction mix was prepared in an Eppendorf tube by adding 10 µl of reconstituted rDNase to 90 µl Reaction buffer for rDNase, enough for all the samples. The reaction mix was briefly vortexed and stored on ice, and when required, 95µl from it was added to each of the RNA samples. Each of the dissected (∼10 mg) tissue was placed into lysis tubes containing 3 metal beads, 320µl lysis buffer RA1 and 3.5µl TCEP. Using Fast prep −24 homogenizer (MP Biomedicals United Kingdom), tissues were disrupted and homogenized as they were lysed 3 times at speed 5 for 20 seconds. Each time the tissues were lysed, the lysis tubes were removed and placed on ice for 1 minute so as to prevent overheating. The lysate was added to Nucleospin filter unit, which was then placed in a 2ml collection tube. RNA was extracted from the lysate after centrifugation at 13,000 rpm for 1 minute. RNA binding condition was adjusted by adding 320µl of 70% ethanol to each homogenised lysate in the collection tubes after removing the filter units. The content of the collection tubes were properly mixed by pipetting up and down for 5 times, using a pipette man. This was then transferred to a NucleoSpin RNA column in a 2ml collection tube alongside precipitate which had formed. The samples were centrifuged for 30 seconds at 10,00rpm and the flow-through discarded.350 µl of membrane desalting buffer (MDB) was added to the NucleoSpin RNA column and centrifuged at 13,000rpm for 1 minute and then the flow-through was discarded. To digest the DNA, 95µl from the DNase reaction mixture was added directly onto the centre of silica membrane of each of the columns and was then incubated at room temperature for 15 minutes. The silica membrane was washed and dried by first adding 200µl RA2 to each of the spin columns and centrifuging them for 30 seconds at 10,000rpm. This was shortly followed by placing the columns in fresh collecting tubes. Furthermore, 600µl buffer RA3 was added to the columns in the fresh tubes and then centrifuged for 30 seconds at 10,000rpm. The flow-through was discarded and the columns dried completely by further adding 250µl of RA3 buffer and centrifuging it for 2 minutes at 13,000rpm. Buffers RA2 and RA3 are wash buffers. While RA2 was used to inactivate the rDNase, the more concentrated RA3 was used to efficiently wash the inner rim and dry the membrane completely (Macherey-Nagel, 2015). The columns from the collection tubes were removed and transferred to a 1.5ml Eppendorf tube and 60µl RNAse free water was added directly onto the column membrane. The membranes were completely covered and then centrifuged at 13,000rpm for a minute. The columns were discarded and the tube containing the eluted RNA samples kept on ice before being stored at −20 °C till the next day.

To avoid contamination and degradation of the RNA, proper care was ensuring gloves were worn and frequently changed when handling RNA samples. Also, reactions were always prepared and kept on ice, thereby preventing RNA degradation.

### RNA quantification

Quantification of the RNA samples was done by using NanoDrop^®^ ND-1000 UV/V Spectrophotometer software (Thermo Fisher Scientific Inc., Waltham, MA, USA) and then calculating the volume of tissue necessary for reverse transcription. The fibre optic measurement surface of the Nano drop device was first cleansed with alcohol to avoid contamination. This was followed by blanking with 1µl RNAse free water directly dropped onto the optic measurement surface. This was followed by clicking the blank and ensuring the wavelength was set to zero, which was the absorbance background against which the samples were measured. Each sample was measured in duplicate and the readings presented in table 2. To know how much of each sample would be used, calculation was made based on the sample with the lowest concentration (85ng/µl) and the required volume of water was added to normalize the tissues (table 2).

**Table 2.**
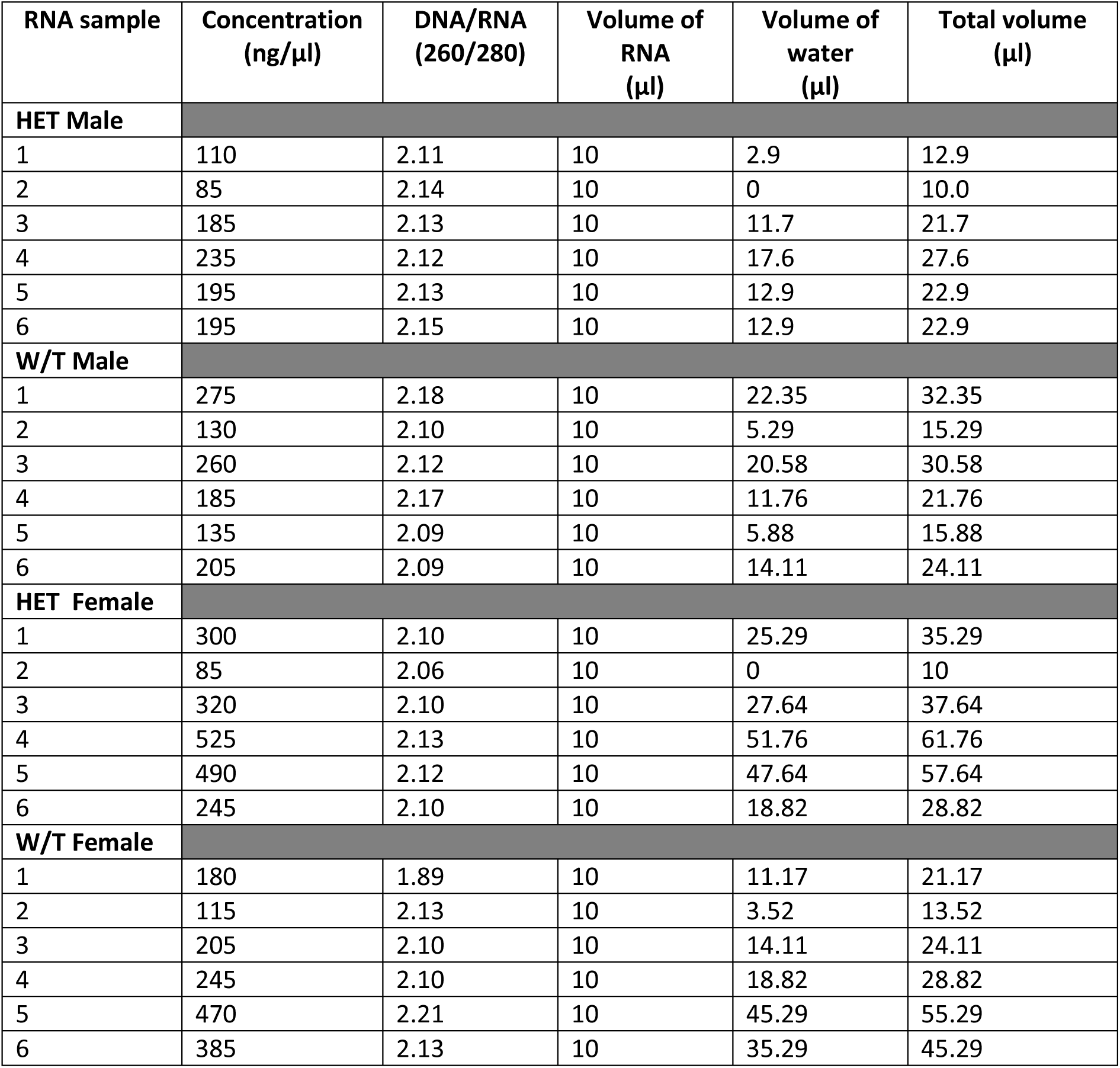
Quantified RNA samples obtained after nanodrop and the volume of water added to each sample to normalise it to 85ng/µl. The concentrations and DNA/RNA ratio are in duplicate values.

| RNA sample | Concentration<br>(ng/μl) | DNA/RNA<br>(260/280) | Volume of<br>RNA<br>(μl) | Volume of<br>water<br>(μl) | Total volume<br>(μl) |
| --- | --- | --- | --- | --- | --- |
| **HET Male** |  |  |  |  |  |
| 1 | 110 | 2.11 | 10 | 2.9 | 12.9 |
| 2 | 85 | 2.14 | 10 | 0 | 10.0 |
| 3 | 185 | 2.13 | 10 | 11.7 | 21.7 |
| 4 | 235 | 2.12 | 10 | 17.6 | 27.6 |
| 5 | 195 | 2.13 | 10 | 12.9 | 22.9 |
| 6 | 195 | 2.15 | 10 | 12.9 | 22.9 |
| **W/T Male** |  |  |  |  |  |
| 1 | 275 | 2.18 | 10 | 22.35 | 32.35 |
| 2 | 130 | 2.10 | 10 | 5.29 | 15.29 |
| 3 | 260 | 2.12 | 10 | 20.58 | 30.58 |
| 4 | 185 | 2.17 | 10 | 11.76 | 21.76 |
| 5 | 135 | 2.09 | 10 | 5.88 | 15.88 |
| 6 | 205 | 2.09 | 10 | 14.11 | 24.11 |
| **HET Female** |  |  |  |  |  |
| 1 | 300 | 2.10 | 10 | 25.29 | 35.29 |
| 2 | 85 | 2.06 | 10 | 0 | 10 |
| 3 | 320 | 2.10 | 10 | 27.64 | 37.64 |
| 4 | 525 | 2.13 | 10 | 51.76 | 61.76 |
| 5 | 490 | 2.12 | 10 | 47.64 | 57.64 |
| 6 | 245 | 2.10 | 10 | 18.82 | 28.82 |
| **W/T Female** |  |  |  |  |  |
| 1 | 180 | 1.89 | 10 | 11.17 | 21.17 |
| 2 | 115 | 2.13 | 10 | 3.52 | 13.52 |
| 3 | 205 | 2.10 | 10 | 14.11 | 24.11 |
| 4 | 245 | 2.10 | 10 | 18.82 | 28.82 |
| 5 | 470 | 2.21 | 10 | 45.29 | 55.29 |
| 6 | 385 | 2.13 | 10 | 35.29 | 45.29 |

### Primer Design – Reverse transcription (RT)

The precision nanoScript 2 RT kit used was supplied by Primerdesign Ltd., Southampton, UK. Frozen reagents were thawed and briefly centrifuged while the rest reagents were placed on ice. 9µl of (85 ng/µl) RNA samples, 0.5 µl Random hexamers and 0.5µl Oligo dTs was added to a thin 0.2ml PCR tube, making up to 10µl. A lid was applied to each sample and was placed in the PCR machine (Biometra GmbH,GÖttingen, Germany) and programmed to heat the samples to 65°C for 5 minutes, which was the annealing temperature (used as per manufacturer’s protocol). After 5 minutes, the samples were retrieved and immediately placed on ice for cooling. An RT master mix was made in a thin walled 0.2 ml PCR tube. For every RT reaction, 10µl of the master mix was added to each sample on ice. The samples were covered with lids and vortexed briefly following a pulse spin and incubated as follows on the PCR machine: 25°C for 5 minutes, 42°C for 20 minutes (extension step), and 72°C for 15 minutes (inactivation step). After the PCR has completed the RT process, the cDNA samples were removed from the machine and stored at −20°C till further use.

### Dilution of cDNA

When required, 3µl each of stored cDNA was transferred into fresh PCR tubes. cDNA was diluted with 27µl RNAse free water, making it up to 30µl. From the 30µl diluted cDNA, 5µl was transferred to a 0.2ml fresh PCR tube. The diluted cDNA was then stored at −20°C for later use.

### Reverse transcription – Polymerase Chain Reaction (end point PCR)

The primers used to amplify the cDNA and their expected product lengths were supplied by Applied Biosystems™ (Roche molecular systems, Inc., Branchburg, USA)and were given as follows: eNOS forward primer: 5’TGGAGAGAGCTTTGCAGCAG’3 and the reverse primer: 5’GATATCTCGGGCAGCAGCTT’3 (424 bp); β-actin forward primer: 5’CCCTGAACCCTAAGGCCA’3 and the reverse primer: 5’AGGATTCCATACCCAAGAAGGAA’3 (489 bp) (figure 2). The eNOS primers were delivered in powdered form as 25 nmoles and were therefore reconstituted with 1000µl RNAse free water to result in final concentration of 25pmol/µl.

**Figure 2.**
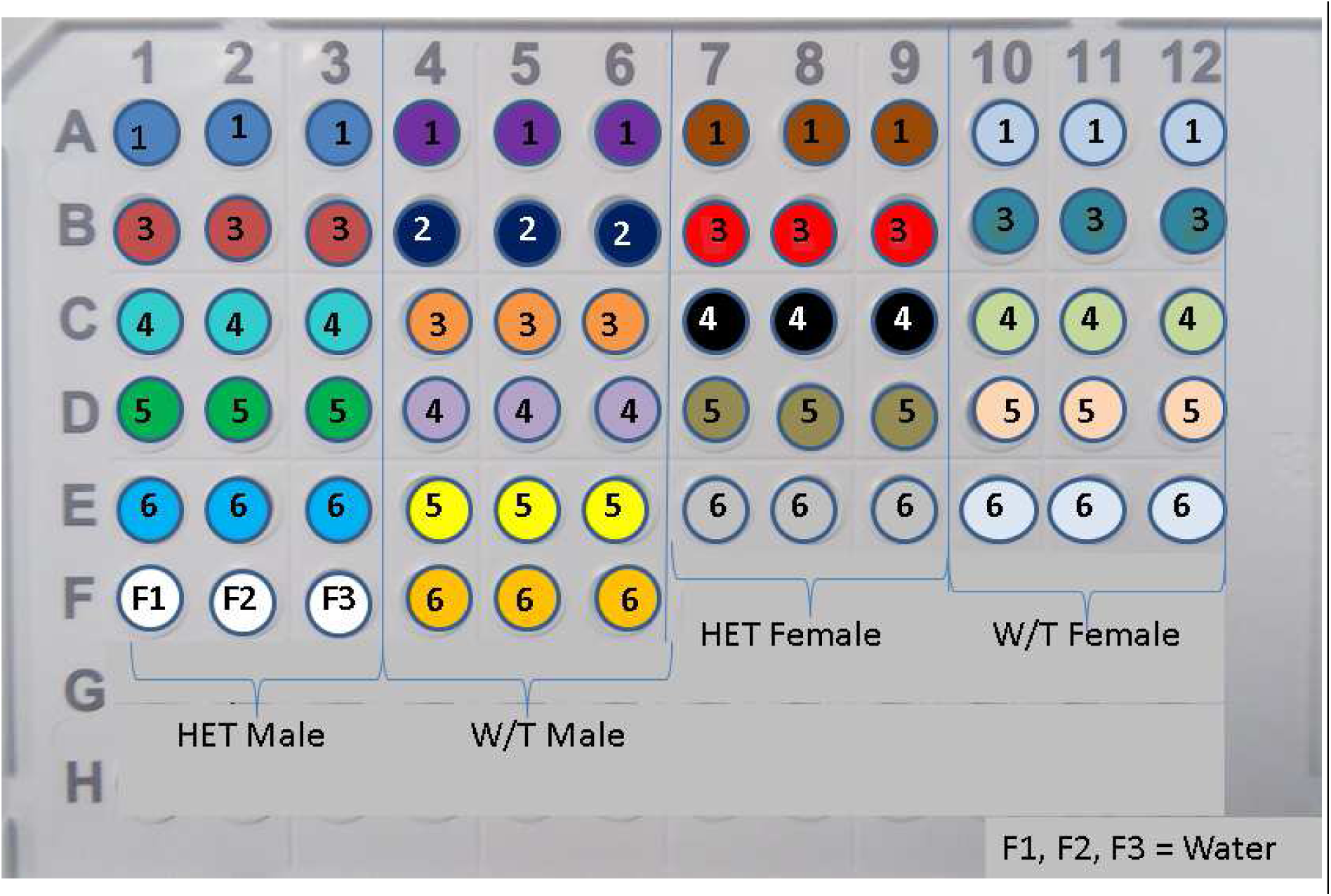
Shows a presentation of how cDNA from 21 lung tissue samples, together with UBC or eNOS (respectively) was introduced into the 96 well qPCR plates in triplicates. Coloured wells and numbers inscribed corresponds to cDNA extracted from any of HET male, W/T male, HET female and W/T female. Wells F1, F2 and F3 are the control wells.

A master mix for all 24 samples containing 25µl Reddy mix, 0.3 µl of forward eNOS primer and 0.3µl of reverse primer was made. The master mix was mixed by vortexing, and 26 µl from it was added to the 5 µl cDNA in the PCR tube. The reagents were mixed and briefly centrifuged to ensure contents were at the bottom of the tube. A PCR negative control was made with 26µl from the master mix and 5µl RNAse free water instead of cDNA template, ensuring that the primers and the ReddyMix were free of DNA contamination. The tubes were transferred to the PCR machine and were programmed at 95°C for 2 minutes (initial denaturation step), 34 x (94°C for 30 seconds (denaturation) /55°C for 60 seconds (annealing step) /72°C for 60 seconds (extension step)) and 72°C for 10 minutes (final extension). The RT-PCR was also carried out for the housekeeper gene, β-Actin with 0.6µl β-Actin added to each cDNA sample. At the end of the PCR, the cDNA samples were retrieved and stored at −20°C.

### Gel electrophoresis

The specificity of the amplified gene and hyperladder was checked with gel electrophoresis analysis on a 1% agarose gel containing 4µlmidori green (Nippon Genetics Europe GmbH, Düren, Germany) and 0.5X TBE (tris / borate / EDTA) solution made from 5X TBE solution following dilution. The 5X TBE was composed of 27g of Tris base powder, 13.5g of powdered boric acid and 1.6ml liquid Ethylenediaminetetraacetic acid (EDTA), all purchased from Fisher Scientific UK Ltd., Loughborough, UK. 10µl of 100bp hyperladder (NorgenBiotek Corp., Thorold, ON, Canada), 20µl of negative control and 20µl each of all cDNA samples incubated with eNOS gene were loaded onto the gel tank and was electrophoresed for 45 minutes at 120 Volts. Electrophoresis was again carried out for β-actin using the process detailed above but with 5µl of 100bp hyperladder IV (Bioline Reagents Limited, London, UK) as it was the only available hyperladder. The gel was photographed using Molecular Imager^®^ Gel Doc™ XR+ System with Image Lab™ Software (Bio-Rad Laboratories Ltd, UK) and the results shown in figure 1.

**Figure 1.**
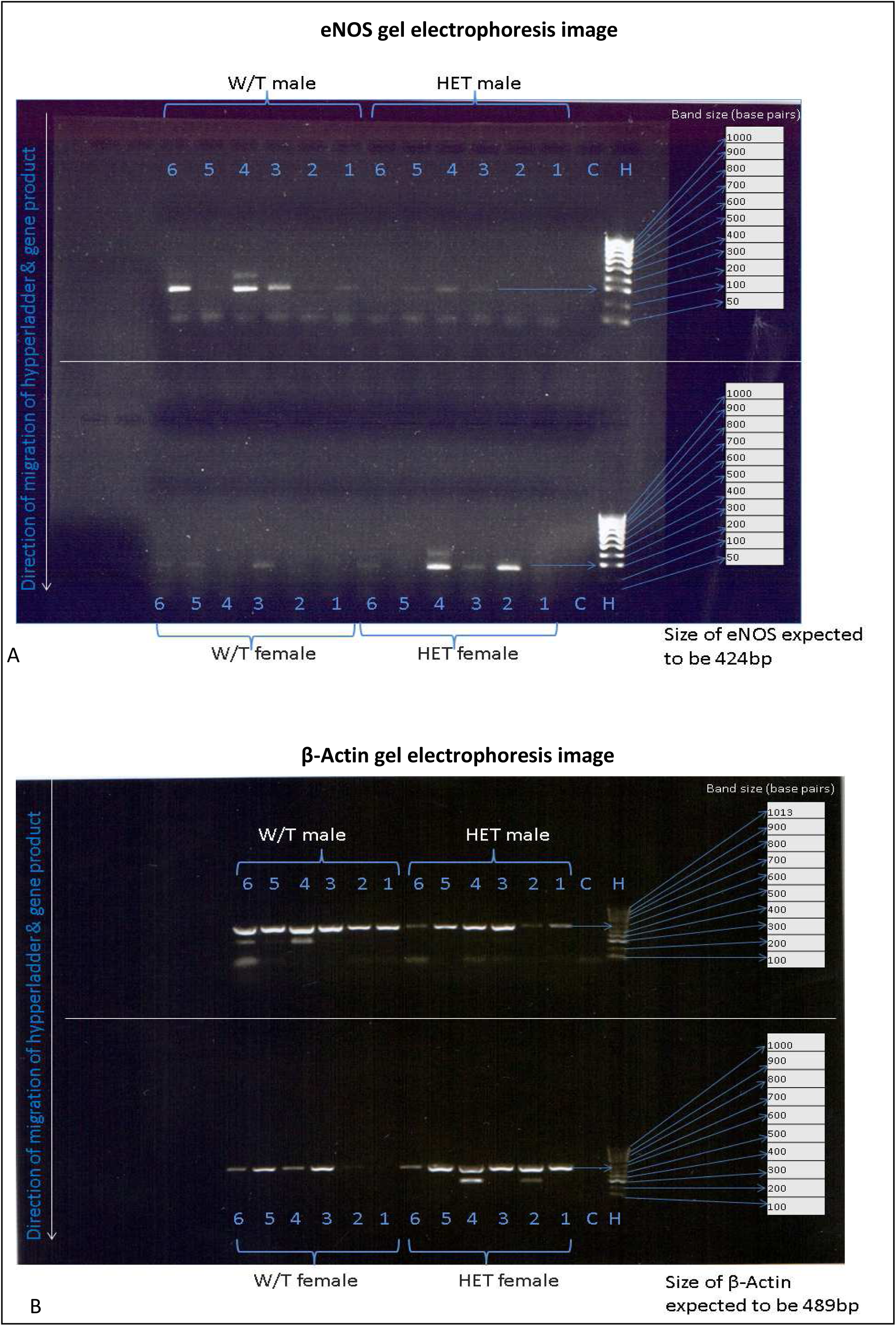
Gel electrophoresis image. Showing expression of eNOS gene (A) and β-Actin (B), control (C),with reference to 100bp hypperladder (H). The numbers represent each sample in a well, corresponding to cDNA from HET male (n = 6), W/T male (n = 6), HET female (n = 6)and W/T female (n = 6). Downward arrows indicate direction of migration of the samples. The expected gene size is also indicated.

### Quantitative Real - Time Polymerase Chain Reaction (qRT-PCR)

The primers used, as well as the 2x qPCR master mix kit were purchased from Primerdesign Ltd., Southampton, UK. First, into fresh PCR tubes on ice, 2µl each of stored cDNA was diluted with 18µl RNAse free water, from which 1.5µl was transferred to a 96 well plate in triplicate for Real-time PCR (figure 2). A PCR negative control without cDNA was also made in triplicate with 1.5µl RNAse free water. A master mix containing 5µl 2x master mix, 0.5 µl eNOS primer and 3µl water was made, sufficient for only 21 samples. The master mix was vortexed and 8.5 µl from it was added to each of the 96 well plate containing the 1.5 µl cDNA. The plate was covered with a film and briefly centrifuged using Allegra X-22R Centrifuge (Beckman Coulter Ltd., High Wycombe, UK) to ensure contents were at the bottom of the plate. The plate was thereafter placed in the ViiA™ 7 Real-Time PCR System (Life Technologies Ltd, Paisley, UK) and was programmed to cycle at 50 °C for 2 minutes, 95 °C for 10 minutes and 40 x (95 °C for 15 seconds/ 60 °C for 1 minute). The 50 degrees step was required to activate the uracil N-glycosylase (UNG) and to remove any contamination. The 95 degree step for 10 minutes was the Taq activating step, at same time denaturing the UNG. At 95 °C for 15 seconds, the dsDNA template denatured. The annealing and extension of the primers by the Taq occurred at 60 °C for 1 minute [33]. Real-time PCR was also done using the housekeeper, ubiquitin C (UBC) primer. The different templates of the gene served as coding and non-coding strands, with only the non - coding strand being transcribed during PCR as it contains the anti-codons [34]. The 2 x qPCR master-mix used was labelled with ROX™fluorescent dye. The qPCR links the amplification of DNA to the generation of fluorescence of ROX dye and was detected during each PCR cycle in real time. The higher the amplified DNA, the higher the intensity of the fluorescence [35]. At the end of the qPCR, the data was first analysed on Microsoft office Excel 2010 (Microsoft Corporation, Redmond, USA). The quantitation cycle (CT) values for all the samples were obtained. The delta delta (ΔΔ) CT method was applied in calculating the change in gene expression. For example, for HET male 1, the average eNOS CT value for the triplicate was 27.665. Average UBC CT value was 24.308. Therefore, ΔCT value for HET male 1: GOI (HET male1) – HK (HET male 1) = 27.665 – 24.308 = 3.357. With the Excel, the average ΔCT values for all HET male were obtained and used in calculating the ΔΔCT values for all other groups. Using GraphPad Prism™ (GraphPad Software Inc., La Jolla, USA), data was collated and relative eNOS gene expression presented on a bar chart while statistical analysis was done using two-way ANOVA (figure 3).

**Figure 3.**
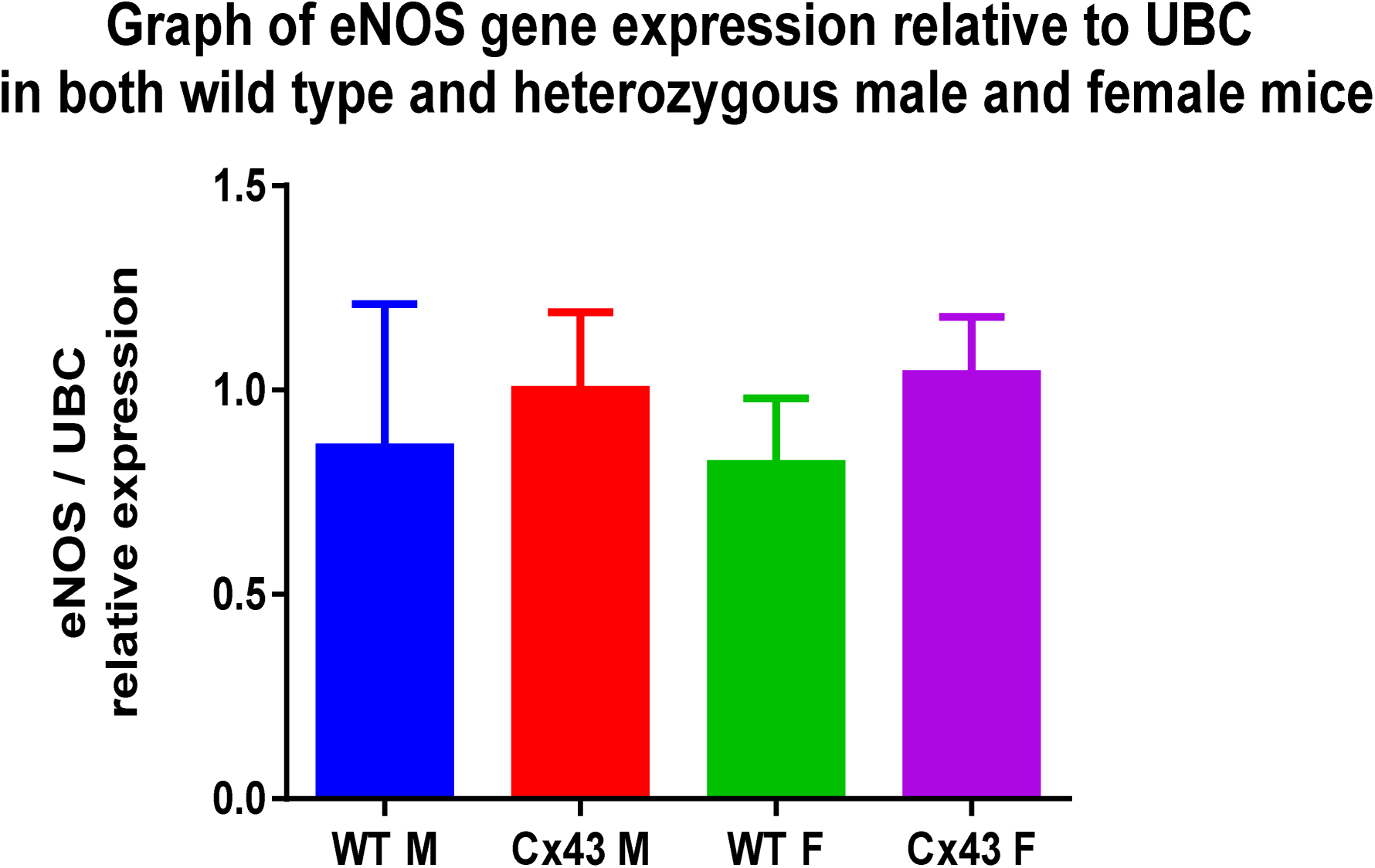
Expression of eNOS. Shows mean values ± SEM of relative gene expression levels measured by qRT – PCR in lung tissues from W/T male (n =6, blue column), HET male (Cx43 M),n = 5, red), W/T female (n = 5, green) and HET female (Cx43 F, n = 5, pink).

### Statistical analysis

Results are expressed as mean ± standard error of mean (SEM), and n shows the number of mice used for end point PCR and qPCR. For end point PCR, gene expressions were visually analysed by looking at the bands and comparing them with the 100bp hyperladder. For qPCR, eNOS gene expression was analysed using two - way ANOVA followed by multiple comparison tests to compare the gene expression levels in both genotypes and phenotypes. A value for p < 0.05 was considered to be significant.

## RESULTS

### End point PCR

To investigate the gene expression of eNOS in heterozygous Cx 43 mice, end point PCR was used to measure the eNOS gene. PCR analysis was done with eNOS and β-Actin transcripts. PCR analysis of eNOS showed that the only visible eNOS bands were in W/T male and HET female and had band size of 200bp. This was however different from the expected size of 424 bp when compared with the hyper ladder (figure 1A). Though, β-Actin was expressed in most of the samples, W/T female was not visible. The size of β-Actin detected was 600bp which was also different from the expected size of 489bp (figure 1B). There were no bands detected in the negative controls.

### qRT - PCR

For more efficiency and reliability, qRT – PCR was used to check the specificity of amplified gene product. eNOS gene was detected in lung tissues relative to UBC (figure 3). Using two - way ANOVA, it was observed that there was no significant difference in the gene expression of eNOS in all four groups of mice models: HET male (1.0 ± 0.19, n = 5), W/T male (0.86 ± 0.35, n = 6), HET female (1.04 ± 0.14, n = 5) and W/T female (0.82 ± 0.16, n = 5). Both HET (Cx43) male and HET (Cx43) female mice showed approximately equal eNOS gene expression levels when compared with their W/T counterparts. There was no change in levels of expression of eNOS in W/T male and W/T female. When statistical analysis was performed at 95% confidence interval, there was no significant difference in eNOS gene expression in both genders (P = 0.98) and genotype (P = 0.43).

## DISCUSSION

The present study tested the expression of eNOS gene in genetically heterozygous Cx 43 (Cx 43 +/-) mice conducted in four groups of mice: W/T male, Cx43 +/- male, W/T female and Cx43 +/- female. The investigation for eNOS expression from the mice lung tissues was first carried out using RT – PCR. Since the bands do not correspond to the expected size of eNOS gene (434bp), it was suggested that the eNOS primers did not pick up the eNOS gene from the cDNA. Also, the negative control wells did not show any bands as they did not contain any cDNA. The housekeeping genes are essential in cellular maintenance in humans and animals and are thought to maintain constant expression levels in all cells, and so, are used as normalizing standards in gene expression [36]. β-Actin was expressed in most of the samples, however, the few samples which appeared not visible suggests insufficient cDNA or contamination. The discrepancy in the size of β-Actin could be also bedue to primers not picking the gene. The use of agarose gel electrophoresis in measuring gene expression of eNOS was quick and easy to cast, it was however unsuitable in separating low molecular weight samples [37], hence the need for qRT – PCR. UBC was shown to be present in the target DNA under test and was used to confirm that the qRT-PCR process worked properly. Since there was no change in the gene expression levels of eNOS, it was therefore suggested that eNOS expression (and possibly its signalling pathway) is not impaired in mice heterozygous in Cx43 (Cx43 +/- mice). Also, gender does not affect eNOS expression as there was no observed change in its expression in both genders. This observation was however not consistent with a recent study by [38] in which sex-dependent alterations of eNOS protein expression in cardiac tissues of male and female W/T mice with heart failure was investigated. Female W/T mice eNOS protein expression levels and activation was shown to be higher than W/T males, and the difference could be due to the presence of eNOS dephosphorylating calcineurin-?A found in W/T males [38].

Recent studies have shown Cx43 to be essential in coordinating cell proliferation and migration in vascular walls [39]. This was confirmed in smooth muscle cells of Cx43 gene knockout mice in which manifestation of accelerated growth of the neointima and the adventitia was observed [28]. Diabetes – impelled inhibition of Cx 43 expression adds to vascular cell apoptosis in retinal endothelial cells of diabetic mice [40]. Quantitative evaluation of Cx 43 expression was shown to be significantly reduced in borderline hypertensive rats (P <0.05) and spontaneously hypertensive rats (P <0.05) compared to normotensive controls [41]. The reduced Cx 43 expression was shown to contribute to remodelling of vascular cell wall and also affect gap junction communication, which was more pronounced in fully established hypertensive rats [41]. This suggests that the accelerated growth of the SMC correlates with Cx43 deletion. However, another finding revealed reduced neointimal formation in Cx 43 heterozygous mice [30]. Interestingly, deletion of Cx43 in mice endothelial cells resulted in hypotension [42] as Cx43 expression was shown to be upregulated in pulmonary arteries from chronically hypoxic rats [24]. However, a decrease in the expression of Cx43 was observed in rats’ aorta when nitric oxide synthase was inhibited [43]. Mice with endothelial cell (EC) specific deletion of Cx43 showed severely reduced Ca^2+^ communication between the EC [44]. Since NO biosynthesis in the EC depends on activation by Ca^2+^ - calmodulin complex, and since there has been evidence that mutant mice which lack eNOS gene appears to be hypertensive [20], a link between Cx43 and eNOS expression has been established, though, with yet to be defined pathway [45]. Since vascular connexins do not operate in isolation, modifying the expression of one connexin may affect the expression of the other [46, 42], and could also influence eNOS gene expression.

Given the different effects of Cx43 deletions in animal models, and the expression of eNOS gene in the mice tested, future work is therefore recommended in understanding the pathogenic basis of PAH. Reactive oxygen species, as well as the interaction of caveolin −1 and other Cx with eNOS affects eNOS activity [47], and this should be taken into account when investigating eNOS expression within a larger population of mice models. How these interactions contribute to rise in PAP could then be measured by taking hemodynamic measurements of lung tissues from the mice under investigation.

## CONCLUSION

In conclusion, the findings have demonstrated that eNOS gene is expressed in both W/T mice and mice heterozygous in Cx43 (Cx43 + / - mice). The expression level of eNOS gene has been shown not to be influenced by gender, and that Cx43 is involved in the development of PAH. Even though eNOS mutant mice seem to develop hypertension, the interaction of Cx43 and eNOS signalling pathway in the pulmonary vasculature appears to be a complex one. Based on these evidences, there are many routes for further research into the interaction of eNOS and other connexins, and how these interactions could affect the expression of eNOS which could possibly lead to the development of PAH.

## ACKNOWLEDGEMENT

Firstly I would like to thank my supervisor, Dr Yvonne Dempsie for supervising this project and giving me the opportunity to carry out this research.

Particular thanks to my extended family and siblings, especially Comr. (Sir) Messiah Wilfred and HM (King) & Mrs T. M. Jamala II for their incredible support and words of wisdom.

Let me acknowledge my ever striving parents, Chief & Mrs Wilfred Uriah. I dedicate this research project to you. I could not have got through this challenging time without your support and guidance.

Finally, I give God the glory for his protection, mercies and faithfulness throughout my stay in the United Kingdom.

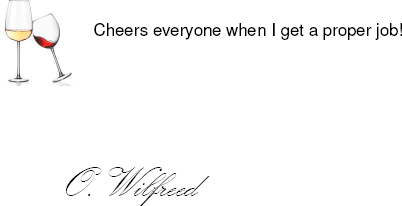

## ABBREVIATIONS

Cx 43: Connexin 43
eNOS: Endothelial nitric oxide synthase
NO: Nitric oxide
PAH: Pulmonary Arterial Hypertension
HET: Heterozygous
W/T: Wild type
RNA: Ribonucleic acid
cDNA: Complementary DNA
PCR: Polymerase chain reaction
PAP: Pulmonary arterial pressure
SMC: Smooth muscle cell
PVR: Pulmonary vascular resistance
RV: Right ventricle
WHO: World Health Organization
IPAH: Idiopathic PAH
HPAH: Heritable PAH
APAH: Associated PAH
PPHN: Persistent pulmonary hypertension of the new born
cGMP: Cyclic guanosine monophosphate
PKG: Protein kinase G
PDE-5: Phosphodiesterase −5
Ca2+: Calcium
NIH: National Institutes of Health
DNAse: Deoxyribonuclease
rDNAse: Recombinant Deoxyribonuclease
TCEP: Tris (2-carboxyethyl) phosphine
rpm: Revolutions per minute
MDB: Membrane Desalting Buffer
RNAse: Ribonuclease
RT: Reverse transcriptase
Oligo-dTs: Deoxy-thymine nucleotides
Bp: Base pairs
RT-PCR: Reverse transcription polymerase chain reaction
TBE: Tris/Borate/EDTA
EDTA: Ethylene-Diamine-Tetraacetic Acid
qRT-PCR: Quantitative Real-Time polymerase chain reaction
UNG: Uracil N-glycosylase (UNG)
UBC: Ubiquitin C
Ct: Quantitation cycle
ΔΔ Ct: Delta delta Ct
GOI: Gene of interest
HK: House keeping
ANOVA: Analysis of variance
SEM: Standard error of mean
EC: Endothelial cell

